# Homeostatic control of meiotic G2/prophase checkpoint function by Pch2 and Hop1

**DOI:** 10.1101/2020.04.24.059642

**Authors:** Vivek B. Raina, Gerben Vader

## Abstract

Checkpoints cascades coordinate cell cycle progression with essential chromosomal processes. During meiotic G2/prophase, recombination and chromosome synapsis are monitored by what are considered distinct checkpoints [1–3]. In budding yeast, the AAA+ ATPase Pch2 is thought to specifically promote cell cycle delay in response to synapsis defects [4–6]. However, unperturbed *pch2Δ* cells are delayed in meiotic G2/prophase [6], suggesting paradoxical roles for Pch2 in cell cycle progression. Here, we provide insight into the checkpoint roles of Pch2 and its connection to Hop1, a HORMA domain-containing client protein. Contrary to current understanding, we find that the Pch2-Hop1 module is crucial for checkpoint function in response to both recombination and synapsis defects, thus revealing a shared meiotic checkpoint cascade. Meiotic checkpoint responses are transduced by DNA break-dependent phosphorylation of Hop1 [7, 8]. Based on our data and on the effect of Pch2 on HORMA topology [9–11], we propose that Pch2 promotes checkpoint proficiency by catalyzing the availability of signaling-competent Hop1. Conversely, we demonstrate that Pch2 can act as a checkpoint silencer, also in the face of persistent DNA repair defects. We establish a framework in which Pch2 and Hop1 form a homeostatic module that governs general meiotic checkpoint function. We show that this module can - depending on the cellular context - fuel or extinguish meiotic checkpoint function, which explains the contradictory roles of Pch2 in cell cycle control. Within the meiotic checkpoint, the Pch2-Hop1 module thus operates analogous to the Pch2/TRIP13-Mad2 module in the spindle assembly checkpoint that monitors chromosome segregation [12–16].

## Results and discussion

### Feedback regulation between Hop1, Zip1 and Pch2 influences SC assembly

In G2/prophase of the meiotic program, double strand break (DSB) formation recombinational repair, synapsis of homologous chromosomes is integrated with cell cycle progression [17]. In budding yeast, Zip1-mediated Synaptonemal Complex (SC) assembly promotes cell cycle progression by regulating chromosome-based checkpoint activity [18–20]. The chromosome axis, of which Hop1 is a key component, forms the structural foundation for recruitment of the SC [21]. The functionality of meiotic checkpoints that monitor recombination and synapsis is linked to successful recombination and SC establishment, and the function of Pch2 is associated with both Hop1 and Zip1: Hop1 is a client of Pch2 [22–25], and chromosomal association of Pch2 is tightly linked to Zip1/SC establishment [5, 20, 23, 26–29]. To understand the wiring between these numerous crucial events during meiotic G2/prophase and the role of Pch2 herein, we initially investigated SC formation and its relation to Pch2, Hop1 and Zip1.

SC components, like Zip1, can form extrachromosomal aggregates known as polycomplexes (PCs) [4]. Cells lacking *NDT80* (*ndt80Δ*), a transcription factor required for G2/prophase exit [30, 31], arrest after DSB formation and crossover repair takes place up to the double Holiday junction stage [32]. We observed that PCs form extensively in *ndt80Δ* cells, but co-deletion of Pch2 led to a striking reduction in the frequency of cells displaying Zip1 polycomplexes (Figure 1A and B, and Supplementary Figure 1A-E). The abundance of PCs was unexpected because PCs are generally associated with defects in DSB processing [18]. However, no considerable defects in SC appearance and assembly were observed in cells lacking Pch2, regardless of whether these cells had developed PCs (Supplementary Figure 1B,C and F), **[4, 5]**. The decrease in PCs in *pch2Δ* cells was also counterintuitive because DSB repair is delayed in *pch2Δ* cells [3, 6, 23, 33]. Hence in this case, if PC formation was a result of DSB defect one would have expected more PCs in *pch2.* In fact, PCs were mostly observed in cells displaying extended levels of Zip1 polymerization (i.e Class III (Supplementary Figure 1C)). Quantification of PCs was only performed in cells showing Class III polymerization of Zip1. In line with a role of Pch2 in regulating PC formation independently of DSB defects, loss of Pch2 did not alleviate the pathological accumulation of PCs observed in cells that fail to generate meiotic DSBs [18] (*i.e.* in *spo11-Y135F* cells; Supplementary Figure 1G and H). In *ndt80Δ*, the cellular levels of Zip1 continue to rise (Supplementary Figure 1A), correlating with the increase in the total number of cells showing PCs (Figure 1B). This suggests that PC formation can be interpreted as a proxy of excess Zip1 remaining after successful establishment of SC. Indeed, overexpression of Zip1 leads to abundant PC formation [34]. These data suggest that Pch2 activity might control the total amount of Zip1 that can be deposited on chromosomes, as such indirectly leading to an observed decrease in PC abundance. In line with this hypothesis, we observed a significantly increased association of Zip1 on the chromosomes in *pch2Δ*, without an obvious effect on total Zip1 levels (Figure 1C and Supplementary Figure 1A) suggesting a role of Pch2 in dynamically modulating SC establishment.

**Figure 1.**
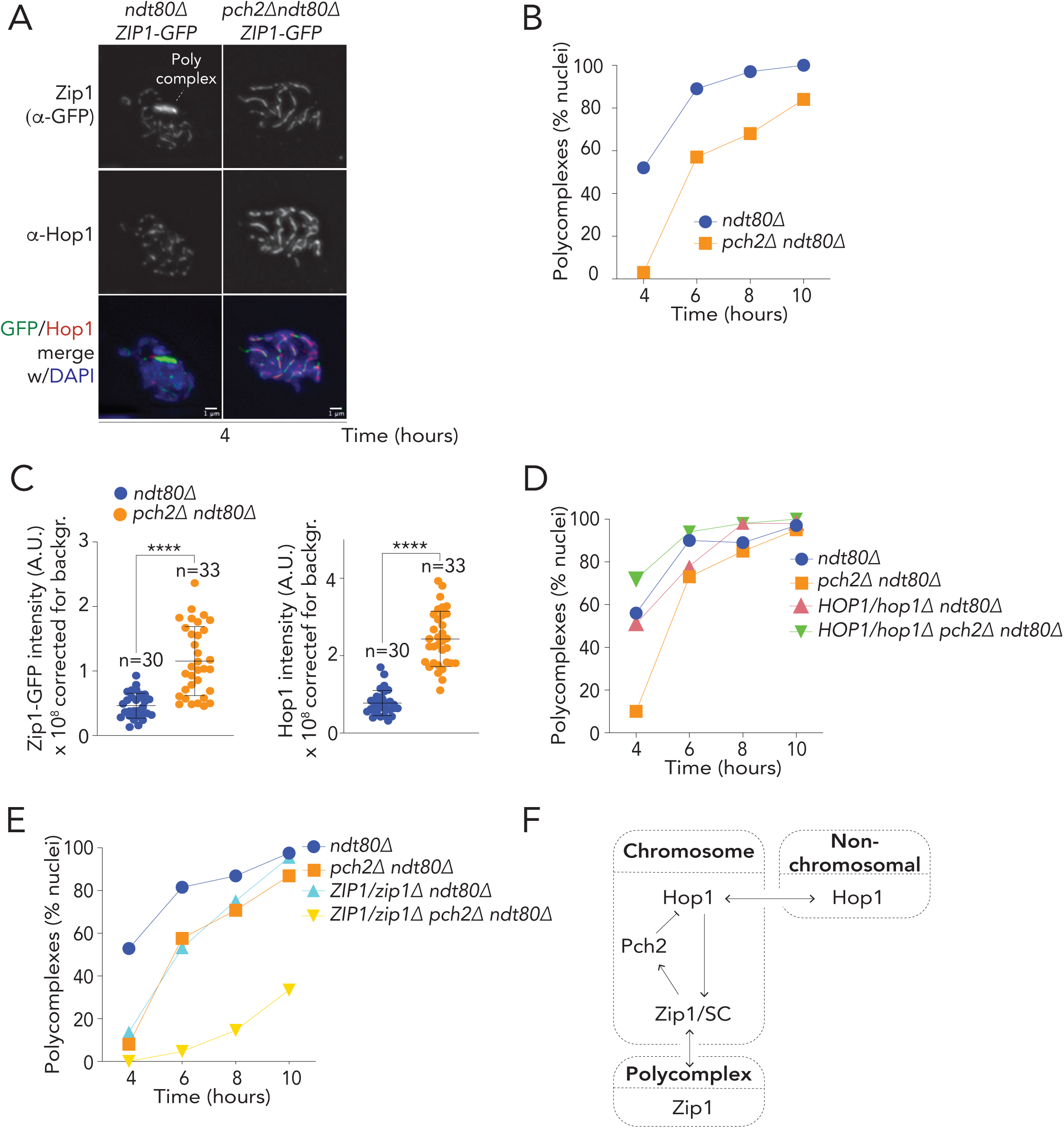
Feedback regulation between Hop1, Zip1 and Pch2 influences SC assembly. **A.** Representative images for Zip1-GFP (green) immunofluorescence and Hop1 (red) immunofluorescence of spread meiotic chromosomes from *ndt80Δ* (yGV4463) and *pch2 ndt80* (yGV4504) strains 4 hours post induction of meiosis. DAPI (blue) was used to stain the DNA. Dotted line shows the polycomplex (extrachromosomal Zip1 aggregate). Scale bar is 1 μm. **B.** Quantification of the number of nuclei showing polycomplexes at indicated time points from same strains as in **A**. Polycomplexes were counted only in cells showing extended polymerization of Zip1 (Class III). A minimum of 100 Class III nuclei were analyzed. **C.** Quantification of chromosomal intensity of Zip1-GFP and Hop1 on meiotic chromosome spreads of strains used in (A) and (B) 4 hours post induction of meiosis. Signal not overlapping with DAPI was excluded and total sum intensity was corrected for background (see Material and methods). The horizontal black line indicates mean and error bars represent standard deviation. **** indicates a significance of *p*0.0001, Mann-Whitney U test. **D.** Comparison of the percentage of class III nuclei, containing polycomplexes in *ndt80Δ* (yGV4463), *pch2 ndt80* (yGV4504), *ZIP1-GFP/zip1 ndt80* (yGV4609), and *ZIP1-GFP/zip1 pch2 ndt80* (yGV4608) strains. At least 100 nuclei were analyzed. **E.** Comparison of the percentage of class III nuclei, containing polycomplexes in *ndt80Δ* (yGV4463), *pch2 ndt80* (yGV4504), *HOP1/hop1 ndt80* (yGV4461) and *HOP1/hop1 pch2 ndt80* (yGV4505) strains. At least 100 nuclei were analyzed. **F.** Schematic showing the feedback regulation between Pch2, Zip1 and Hop1. Arrows indicate functional relationship between factors. Absence of Pch2 increases Hop1 levels on chromosome thus providing more availability for Zip1 to polymerize and hence delay polycomplex formation.

In both yeast and mammals, SC polymerization relies on the Hop1-containing chromosome axis, whereas Pch2-dependent removal of Hop1 (or its mammalian homologs) from chromosomes depends on SC-driven chromosomal recruitment of Pch2 [5, 20, 23, 26–29]. We detected an increased association of Hop1 on meiotic chromosomes in *pch2Δ* cells (Figure 1C) [20, 23]. This increased association of Hop1 correlated with the increase of Zip1 on chromosomes. It is to note that we also observed an increase in the total cellular Hop1 in *pch2Δ* cells (Supplementary Figure 1A) [20], suggesting that the reduced prevalence of PCs in *pch2∆* cells might reflect higher chromosomal levels of Zip1, driven by increased Hop1 levels. We reasoned that changing the cellular levels of Hop1 would be expected to reduce the fraction of Zip1 on chromosomes and increase PC abundance. On the other hand, reducing Zip1 cellular levels would result in decrease in excessive Zip1 being available for PC formation. Indeed, reducing Hop1 levels (by making use of a heterozygous deletion) increased PC abundance in *pch2Δ* cells (Figure 1D, and Supplementary Figure 2A and C) and reducing Zip1 levels (Supplementary Figure 2B and D), led to a reduction in PC abundance (Figure 1E). The reduction in PCs in cells with reduced Zip1 levels was synergistic with *pch2*, likely reflecting a reduced pool of excessive Zip1 protein available for PC assembly. These findings reveal a dynamic Pch2-mediated feedback regulation of SC assembly with chromosomally-localized Hop1 (Figure 1F). We suggest that this Pch2-mediated regulation of chromosomally localized Hop1 influences cellular and chromosomal behavior during meiotic G2/prophase.

### Pch2 and Hop1 collaborate to mediate G2/prophase checkpoint function

Cells lacking Pch2 show increased amounts of chromosomal Hop1 (Figure 1C [20]), and are delayed in progression through meiotic G2/prophase [6]. In cells that lack Zip1, where chromosomal recruitment of Pch2 to remove Hop1 is impaired [5], removal of Pch2 contrastingly leads to accelerated cell cycle progression [4, 5]. However, *pch2Δ* does not affect cell cycle arrest due to lack of Dmc1 [6], a meiosis-specific RecA homolog crucial for meiotic homologous recombination [35–37]. Thus, under different conditions (*i.e.* in wild type, *zip1Δ or dmc1Δ cells*), Pch2 contributes differently to cell cycle control. We aimed to address the molecular basis for these apparent distinct phenotypic behaviors of *pch2Δ* cells. Inspired by our observations revealing a functional connection between Pch2 activity, Hop1 protein levels and SC assembly (see above and Figure 1), we investigated if Hop1 abundance affected these Pch2-associated phenotypes. We queried progression through meiotic G2/prophase by analyzing entry into Meiosis I/II (as scored by spindle morphology) and by activation of Ndt80, which we traced by monitoring expression of Cdc5, a key transcriptional target of Ndt80 [31] (Figure 2A). Interestingly, we observed that the *pch2Δ-* dependent delay was alleviated by reducing the levels of Hop1, underscoring our earlier observation that Hop1 levels influence Pch2-associated phenotypes (Figure 2B and Supplementary Figure 3A). The effects on cell cycle progression were associated with effects on checkpoint activity which was analyzed by monitoring phospho-status of Hop1 via well-documented electrophoretic mobility shift (Hop1 is phosphorylated by Mec1/Tel1 kinases in response to checkpoint-activating conditions [7, 38]), and activation of Mek1, a G2/prophase checkpoint kinase, by querying the phospho-status of Histone H3-threonine (pH3-T11) (a confirmed Mek1 substrate) [39] (Figure 2A). We also observed similar effects between Pch2 and Hop1 when analyzing spore viability (Supplementary Figure 3B). These findings show that Hop1 protein levels influence Pch2-associated phenotypes beyond SC assembly.

**Figure 2.**
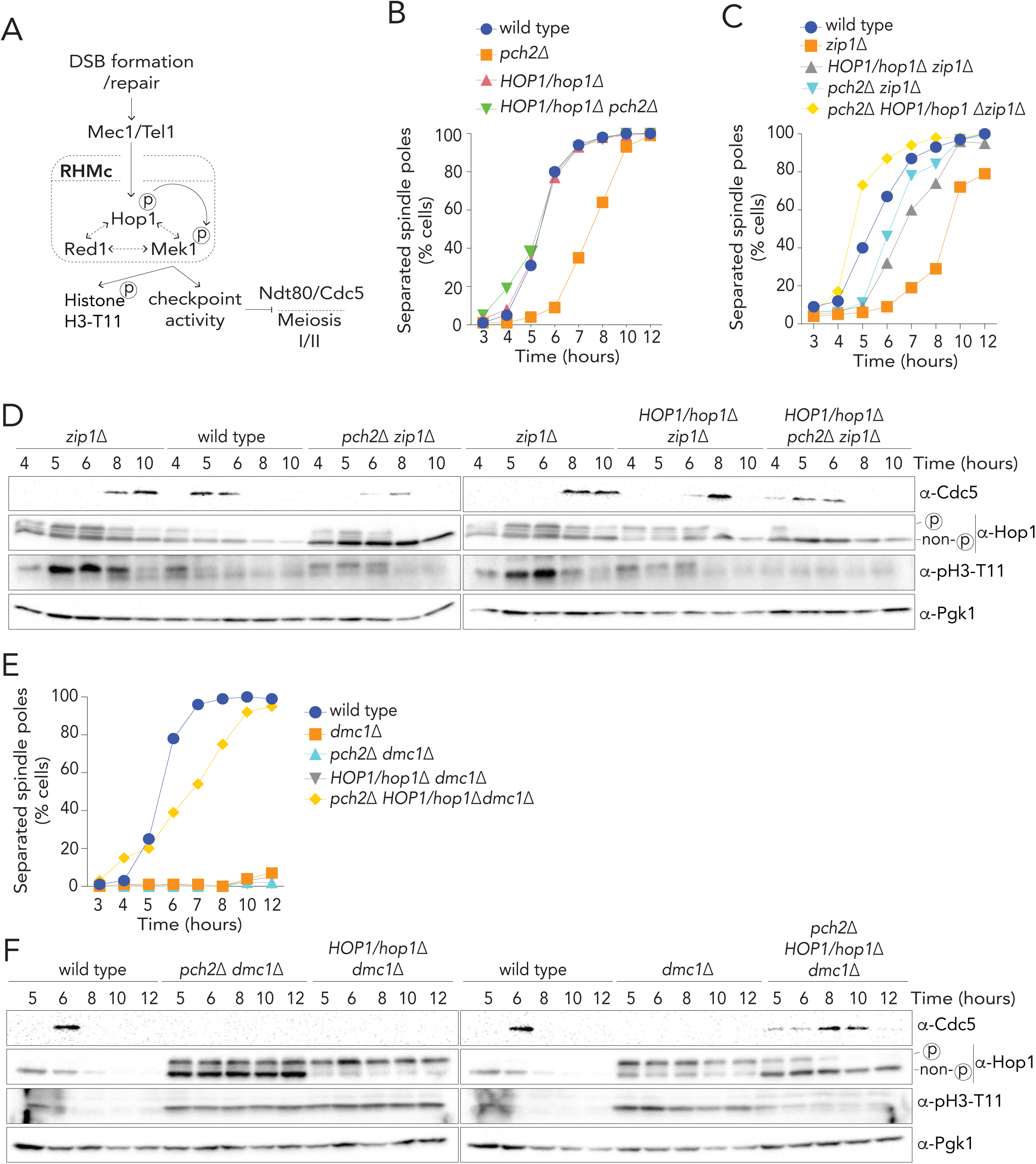
Pch2 and Hop1 collaborate to mediate G2/prophase checkpoint function. **A.** Schematic showing proteins involved in signaling cascade of DSB repair and progression and proteins involved in Meiosis I entry. **B.** Meiotic cell cycle progression analyzed by -tubulin immunofluorescence to count percentage of cells showing separated spindle pole in wild type (yGV4527), *pch2Δ* (yGV4528), *HOP1/hop1* (yGV4526) and *HOP1/hop1 pch2* (yGV4525). A minimum of 200 cells were counted for each data point. **C.** Meiotic cell cycle progression analyzed by -tubulin immunofluorescence to count percentage of cells showing separated spindle pole in wild type (yGV4577), *zip1* (yGV4578), *HOP1/hop1 zip1* (yGV4579), *pch2 zip1* (yGV4581), *pch2 HOP1/hop1 zip1* (yGV4580) strains at indicated time points. A minimum of 200 cells were counted for each data point. **D.** Western blot analysis of Cdc5, Hop1, phospho-Histone 3-Thr11(pH3-T11) and Pgk1 (loading control) from the same cultures analyzed in **C**. Cdc5 is used as a marker for entry into meiosis I; the slow migrating band of Hop1 (indicated) depicts documented phospho-shift of Hop1, a known marker for Mec1/Tel1 activity; pH3-T11 is used as a marker for Mek1 activity and Pgk1 is used as a loading control. All the samples are from the same time course experiment. **E.** Percentage of cells showing separated spindle poles as measured by -tubulin immunofluorescence of wild type (yGV4577), *dmc1* (yGV48), *pch2 dmc1* (yGV1269), *HOP1/hop1 dmc1* (yGV4546), *pch2 HOP1/hop1 dmc1* (yGV1171) at indicated time points. A minimum of 200 cells were counted for each data point. **F.** Western blot analysis of Cdc5, Hop1, pH3-T11 and Pgk1 from the same cultures as analyzed in D. All the samples are from the same time course experiment.

Cell that lack *ZIP1* are delayed in G2/prophase with an active checkpoint [5, 35] as indicated by elevated phospho-Hop1 levels, active Mek1 signaling, and delayed Cdc5 expression (Figure 2C and D). Intriguingly, *HOP1* heterozygosity partially alleviated this delay, to an extent that was similar to that observed in *zip1Δ pch2Δ* cells (Figure 2C and D). Thus, optimal synapsis checkpoint function requires the correct cellular levels of Hop1. Combining *pch2Δ* with *HOP1* heterozygosity led to a combined effect on checkpoint dysfunction (Figure 2C and D) showing Pch2 is necessary to fuel the checkpoint under reduced Hop1 levels. This relation between Pch2 and Hop1 was reflected in spore viability defects (Supplementary Figure 3C). We also investigated the requirements for Pch2 and Hop1 levels in a *zip1Δ rad51Δ* background. In this combined synapsis and DNA repair-impaired background, we observed a similar synergistic functional impairment between Pch2 deletion and *HOP1* heterozygosity (Supplementary Figure 3D and E). Our observations indicate that an interplay between Pch2 and Hop1 is required for an efficient G2/prophase checkpoint response to synapsis defects.

Cells that lack Dmc1 show a near-complete arrest in G2/prophase [6, 35], with unrepaired DSBs and persistent Mec1/Tel1 and Mek1 activation. Previous work suggested that Pch2 is not required for checkpoint function in response to recombination defects, leading to the idea that two distinct checkpoint responses monitor meiotic G2/prophase, with the checkpoint function of Pch2 being limited to synapsis defects [6, 40]. Indeed, neither *pch2Δ* nor *HOP1* heterozygosity were able to alleviate the *dmc1Δ*-dependent arrest (Figure 2E and F) [3, 6]. However, the combination of *pch2Δ* and *HOP1* heterozygosity led to the unexpected robust elimination of checkpoint function in cells lacking Dmc1 (Figure 2E and F). This indicates that Pch2 acts synergistically with Hop1 in mediating the recombination checkpoint. Taken together with our observations on the *zip1Δ* checkpoint response, these data argue that Pch2-Hop1 is a central signalling module that is used by both the synapsis defect- and recombination defect-sensing cascades. In *zip1Δ* cells, Pch2 is not associated with chromosomes, except with nucleolar chromatin [5]. The nucleolar pool of Pch2 is unlikely to be involved in checkpoint function [41], and work in *Drosophila* suggests that Pch2 is associated with nuclear lamina [41, 42]. In line with previous observations [43], Pch2 was required for phosphorylation of Hop1 in checkpoint activating conditions: we observed that the abundance of Hop1 and presence of Pch2 affected the phosphorylated pool of Hop1 (for example, compare wild type, *pch2 dmc1, HOP1/hop1 dmc1* and *pch2 HOP1/hop1 dmc1* in Figure 2F). Combined with our findings (see also above; Figure 1) and earlier work [41], these observations suggest a dynamic exchange between non-chromosomal and chromosomal (*i.e.* active) Hop1, driven by (non-chromosomal) Pch2 function influence checkpoint function.

How can non-chromosomal Pch2 influence the checkpoint function of its client Hop1 on chromosomes? We reasoned that the function of Pch2 may lie in maintaining the levels of ‘signaling-competent’ Hop1 required for its dynamic incorporation into ‘checkpoint-active’ chromosomal sites. This hypothesis is inspired by the biochemical characteristics of Hop1, and by the function of mammalian Pch2 homologs (referred to as Pch2/TRIP13 [24]) in spindle assembly checkpoint (SAC) signaling that monitors chromosome segregation.

Hop1 contains a HORMA domain [44], a structurally-conserved domain that can exist in an open/unbuckled or closed confirmation [10, 11, 22]. Closed HORMA domains can topologically embrace proteins by engaging with closure motifs (CMs) present in interacting proteins in a ‘safety belt’ binding mechanism [22, 45]. CM-mediated engagement with a binding partner is thought of as the active state of HORMA-domain containing proteins. In the case of Hop1, CM-mediated interactions promote its incorporation into the meiotic chromosome axis [11, 46–48] and mediates formation of the Red1-Hop1-Mek1 complex (RHMc). When on chromosomes, Hop1 can be phosphorylated by DSB-responsive Mec1/Tel1 kinases [7, 49], which triggers localized activation of Mek1 and downstream checkpoint activity [7, 38, 50, 51]. Pch2, via a direct AAA+ ATPase-client relationship, influences the topology of HORMA domains, catalyzing their conversion from the closed to the open form [9, 22, 24, 52].

A key step in SAC signaling is the generation of the mitotic checkpoint complex (MCC) [53–56], whose formation depends on the CM-based interaction of closed (C-)Mad2 with Cdc20 [57, 58]. Mad2 is a HORMA domain protein that exists in open (O-) and closed states [45, 59], and Pch2/TRIP13 catalytically counteracts the spontaneous conversion of O-Mad2 into C-Mad2 [9, 22, 52]. In response to improper chromosome-microtubule interactions [45, 57–59], O-Mad2 is recruited to kinetochores, which catalyze its incorporation into MCC [53, 60–62]. Failure to maintain a sufficiently large pool of O-Mad2 in cells lacking Pch2/TRIP13 impairs checkpoint function [12, 63]. We suggest that the wiring of meiotic checkpoint signaling is conceptually and biochemically analogous: Pch2 activity is needed to continuously generate an ‘open’-Hop1 molecule– through topological conversion of C-Hop1 –, and this ‘open’ Hop1 is competent to be incorporated into chromosome-based checkpoint catalysis. (Note that the ‘open’ topology of Hop1 is structurally slightly different from the classical O-Mad2 topology, and keeping with the current nomenclature, we will refer to this state of Hop1 as ‘unbuckled’ (U-)Hop1 [11]). This model should be extended to explain why, under certain conditions, Pch2 is redundant in mounting a checkpoint response (*e.g.* in *dmc1Δ*, if Hop1 levels are not interfered with - see above). We postulate that a sufficient concentration of the open form of Hop1 (dictated by the equilibrium between U-Hop1 and C-Hop1, and Hop1 total protein levels), proficient to be incorporated in RHMc, exists even without Pch2. Evidently, in the absence of Pch2 activity, such a system can only generate productive signaling once (pathologically) high levels of Mec1/Tel1-based phospho-signaling are present (as is the case in *dmc1Δ* cells, which experience excessive ssDNA tracts due to ongoing DNA end resection [6, 7, 35]), and when normal levels of Hop1 are maintained (as shown here). Under less extreme kinase signaling conditions (*e.g.* in wild type or *zip1Δ* cells) or when Hop1 levels are compromised, Pch2 becomes crucially important to catalyze the establishment of a large enough pool of checkpoint-competent Hop1. We note that in *pch2*, total cellular Hop1 levels are increased (*e.g.* Supplementary Figure 1A). This might affect checkpoint functionality under specific conditions, by influencing the absolute number of U-Hop1 molecules present within the cell.

### The Pch2-Hop1 module drives meiotic checkpoint silencing

The ability of Pch2/TRIP13 to catalyze the transition of C-Mad2 to O-Mad2 is not only required to fuel kinetochore-driven MCC assembly; it can also lead to the disassembly of MCC complexes once proper chromosome-microtubule interactions are established [9, 13–16, 52]. Thus, in SAC-signaling Pch2/TRIP13 can also promote SAC silencing and cell cycle progression. If the role of Pch2 in regulating meiotic checkpoint function is analogous to the function of Pch2/TRIP13 in SAC signaling, it should thus also exhibit similar checkpoint antagonizing activities during meiotic G2/prophase. There are some observations pointing in this direction. For example, Pch2 can extinguish Mek1-dependent signaling [20], in a manner that is linked to SC-driven chromosomal recruitment of Pch2. In addition, cells that lack Pch2 are delayed in meiotic G2/prophase under conditions of sustained Hop1 phosphorylation (Figure 2B and Supplementary Figure 3A), which ostensibly is caused by impaired disruption of activated RHMc-dependent signaling. We aimed to dissect this silencing function of Pch2 during meiotic G2/prophase. To do so, we first established an inducible Pch2-expression system (based on the established *pGAL*/*GAL4-ER* system [64]) (Figure 3A). Early expression of Pch2 phenocopied *pch2Δ* (in terms of Hop1 phospho-status) (Supplementary Figure 4A and B), and did not lead to spore viability defects (Supplementary Figure 4C), indicating that this system leads to expression of functional Pch2. Timed expression of Pch2 in *ndt80Δ* cells increased PC abundance, mirroring our observations on the relationship between Pch2, Hop1 and Zip1 with regards to chromosomal association and PC formation (see above; Figure 1 and Supplementary Figure 4D-F). This observation also strengthens our earlier deduction that a balance between chromosomal and PC-bound Zip1 is dynamically set by Pch2 and Hop1 (see above and Figure 1). Importantly, induction of Pch2 expression late in meiotic G2/prophase (*i.e.* at t=7 hours), led to a rapid reversal of phenotypes associated with sustained checkpoint function in *pch2Δ* cells (*i.e.* phosphorylated Hop1 and active Mek1) (Figure 3B), indicating that under these conditions, Pch2 induction extinguished checkpoint function during meiotic G2/prophase.

**Figure 3.**
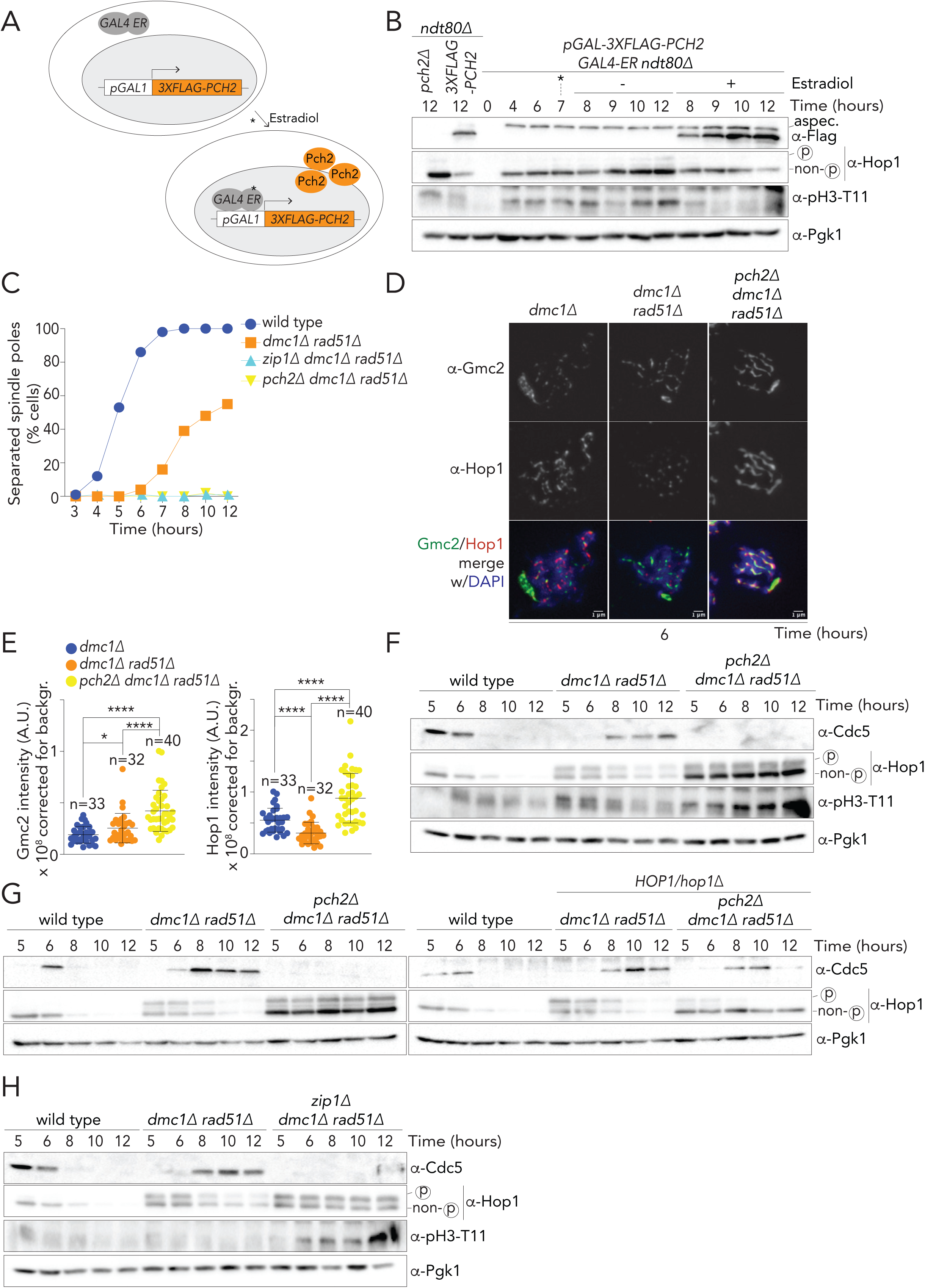
The Pch2-Hop1 module drives meiotic checkpoint silencing. **A.** Schematic showing the estradiol-based *pGAL1*-*3XFLAG-PCH2 GAL4-ER* system. **B**. Western blot analysis of Hop1 and pH3-T11 to compare checkpoint activity with and without induction of Flag-Pch2 in a *ndt80Δ* background (yGV3171). Expression levels are compared with *pch2 ndt80* (yGV2447) and *Flag-Pch2 ndt80* (yGV2889) where Flag-Pch2 is expressed from endogenous promoter of Pch2. Estradiol was added 7 hours post-induction of meiosis. Induction of Pch2 was confirmed by western blot using -Flag antibody. Pgk1 is used as a loading control. **C.** Cell cycle progression analyzed by separated spindle pole of wild type (yGV4577), *dmc1 rad51* (yGV1401), *zip1 dmc1 rad51* (yGV4737) and *pch2 dmc1 rad51* (yGV1409). A minimum of 200 cells were counted for each data point. **D.** Representative images of meiotic chromosome spreads from *dmc1* (yGV48), *dmc1 rad51* (yGV1401) and *pch2 dmc1 rad51* (yGV1409), 6 hours post induction of meiosis and stained for Gmc2 (green) and Hop1 (red). Gmc2 is used to assess SC formation. DAPI (blue) is used to stain DNA. Scale bar is 1m. **E.** Quantification of background corrected chromosomal intensity of Gmc2 and Hop1 on meiotic chromosome spreads from strains shown in **D**. Signal not overlapping with DAPI was excluded. * and **** indicates a significance of *p*0.05 and 0.0001, respectively, Mann-Whitney U test. **F.** Western blot analysis of Cdc5, pH3-T11, Hop1 and Pgk1 (loading control) in wild type (yGV4577), *dmc1 rad51* (yGV1401) and *pch2 dmc1 rad51* (yGV1409). **E.** Western blot analysis to check cell cycle progression and checkpoint activity in Wildtype (yGV4577), *dmc1 rad51* (yGV1401), *pch2 dmc1 rad51* (yGV1409), *HOP1/hop1 dmc1 rad51* (yGV4713) and *pch2 HOP1/hop1 dmc1 rad51* (yGV4715). All the samples are from the same experiment. **G.** Western blot analysis of Cdc5, pH3-T11, Hop1 and Pgk1 (loading control) in wildtype (yGV4577), *dmc1 rad51* (yGV1401) and *zip1 dmc1 rad51* (yGV4737). Results shown in **C**., **F**., and **H**. are from the same time course experiment.

To further expose the checkpoint silencing function of Pch2, we wished to identify a condition in which cells fail to establish an efficient checkpoint response despite high amounts of DSB-based signaling. Cells that lack both RecA-like DNA recombinases required for DSB repair in meiosis (*i.e.* Dmc1 and Rad51) progress past meiotic G2/prophase despite a failure to repair DSBs (Figure 3C) [65, 66]. The reason for this override has remained unclear; in principle a strong arrest in meiotic G2/prophase in response to lack of any recombinase activity would be expected, similar to what is seen in *dmc1Δ.* To shed light on this phenotype, we investigated the chromosomal recruitment of Gmc2, an SC-component [67] in *dmc1Δ rad51Δ* cells. Strikingly, we found that these cells exhibited substantial amounts of chromosomal SC-like structures (as judged by Gmc2-staining), whereas *dmc1Δ* cells contained few SC-like chromosomal structures (Figure 3D and E). This suggests that cells which lack Rad51 and Dmc1 form extensive SC structures in a manner that is uncoupled from DSB repair. Based on this observation, we hypothesized that deleting *RAD51* may rescue the arrest of *dmc1Δ* arrest by allowing (uncoordinated) synapsis [68], enabling recruitment of Pch2 to the chromosomes, and associated checkpoint silencing. Excitingly, we observed that removing Pch2 in the *dmc1Δ rad51Δ* background prevented these cells from prematurely entering Meiosis I/II (Figure 3C and F). In accordance with the model that Pch2-dependent checkpoint silencing is executed via extraction of Hop1 from the chromosome axis [20], we found that the amount of Hop1 on chromosomes was significantly higher in *pch2 dmc1 rad51* as compared to *dmc1Δ rad51Δ* (Figure 3D and E). Thus, even in a recombination-deficient condition (*i.e. dmc1 rad51*) [65, 66], Pch2 can trigger checkpoint silencing. We next explored this condition to investigate how Pch2-dependent checkpoint signaling depended on Hop1. The cell cycle arrest observed in *pch2 dmc1 rad51* did not tolerate reduced Hop1 protein levels (Figure 3G), demonstrating that the Pch2-Hop1 module is crucial for establishing checkpoint function.

Our model explaining Pch2-driven checkpoint silencing predicts that cell cycle progression *dmc1Δ rad51Δ* cells should dependent on SC-formation. In line with this idea, we found that, similar to *pch2*, deleting *ZIP1Δ* prevented checkpoint silencing in the *dmc1Δ rad51Δ* background (Figure 3H). In total, these results reveal that SC-dependent Pch2 function is required for checkpoint silencing, even in the face of unrepaired DSBs. We propose that a key factor that determines whether Pch2 acts as a checkpoint agonist or antagonist is SC-polymerization, and thus Pch2 recruitment to chromosomes. Of note, our results also suggest that cell cycle progression in cells lacking Dmc1 and Rad51 is functionally connected to uncoordinated SC polymerization [65, 66, 68].

To delineate the activation and silencing activities of Pch2-dependent signaling, we exploited our inducible Pch2-expression system (Figure 3A). We aimed to create conditions that either fail to activate or silence checkpoint function in the absence of Pch2, and investigate the effect of Pch2 induction under such conditions. To investigate the role of Pch2 in checkpoint activation, we focused on the *zip1Δ* checkpoint response. We tested the effect of induced expression of Pch2 in a *zip1Δndt80Δ* background, and found that expression of Pch2 triggered checkpoint establishment (Supplementary Figure 4G). Checkpoint activity was already observable, albeit at reduced efficiency, even without the induction of Pch2, presumably because of the presence of sufficient levels of Hop1. This prompted us to investigate the response in a *zip1Δ HOP1/hop1Δ* background, in which loss of Pch2 led to complete checkpoint loss, as shown in Figure 2C. Indeed, under conditions that do not induce Pch2 in *zip1ΔHOP1/hop1Δndt80Δ* cells, we observed a lack of checkpoint signaling (as judged by phospo-Hop1 and activated Mek1) (Figure 4A). Interestingly, upon induction of Pch2, we detected a rapid and robust activation of checkpoint function (Figure 4A). These results argue under condition where Pch2 cannot be recruited to meiotic chromosomes (due to lack of SC formation in *zip1Δ*), expression of Pch2 leads to an activation of the checkpoint, indicating that non-chromosomal Pch2 drives checkpoint activation. Conversely, induced expression of Pch2 in a *dmc1Δ Δrad51* background, led to a rapid decrease of checkpoint-associated markers, and concomitant cell cycle progression into Meiosis I/II, indicative of successful checkpoint silencing (Figure 4B and C). Under these conditions, Pch2 can ostensibly be recruited to chromosomes (due to unscheduled SC formation - see above; Figure 3D and E). Combined with our earlier findings, these data emphasize that depending on the chromosomal context, Pch2 activity can lead to opposite functional outcomes: checkpoint activation or inactivation.

**Figure 4:**
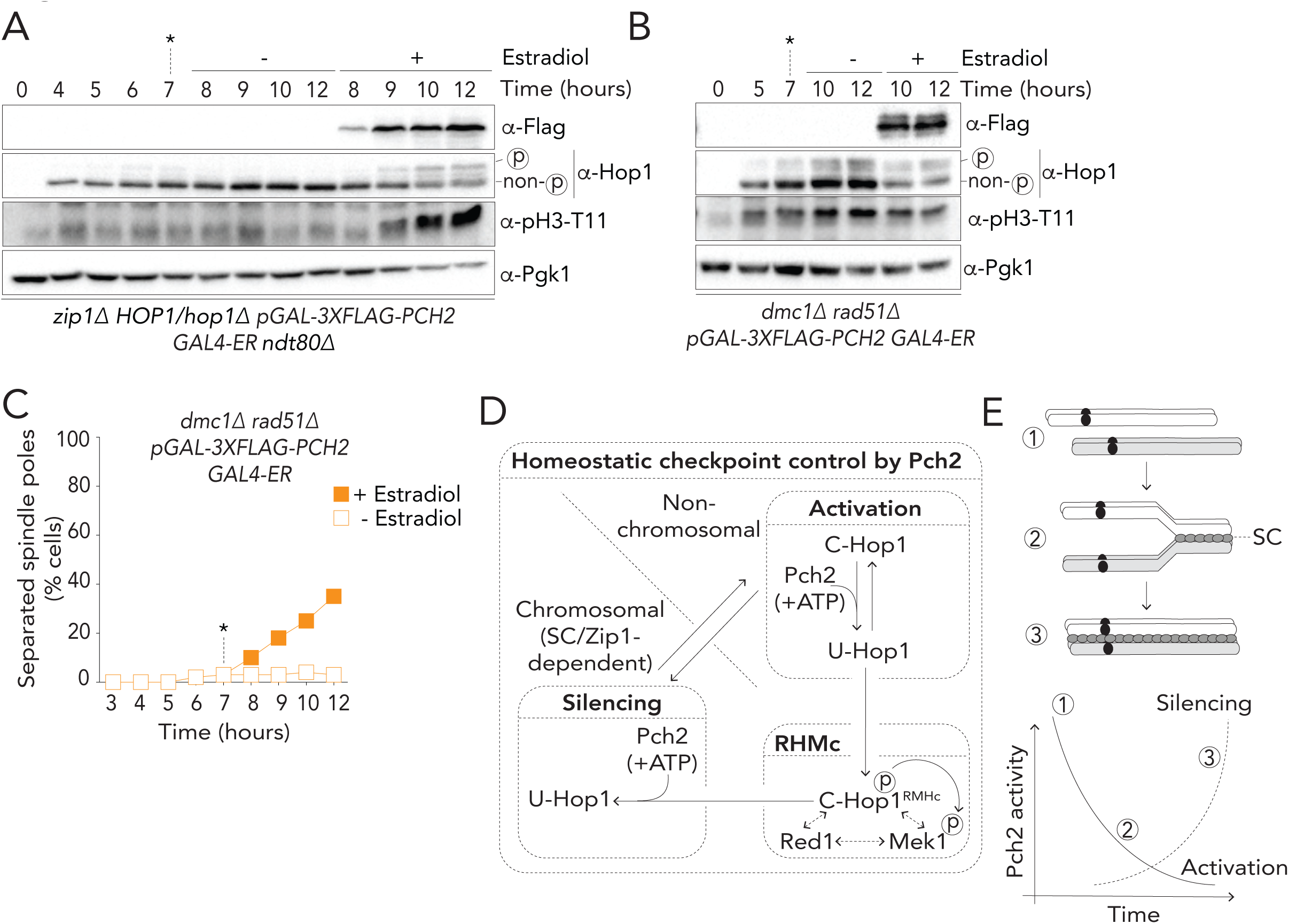
Delineating the dual roles of the Pch2-Hop1 module in meiotic G2/prophase. **A.** Western blot analysis of Hop1 and pH3-T11 with and without induction of Flag-Pch2 in *zip1Δ HOP1/hop1Δ ndt80Δ* (yGV4637). Estradiol was added 7 hours post induction of meiosis (indicated by *) and induction of 3xFlag-Pch2 was confirmed by western blot using -Flag antibody. Pgk1 was used as a loading control **B.** Western blot analysis of Hop1 and pH3-T11 analyzed with and without induction of 3xFlag-Pch2 in *dmc1Δ rad51Δ* (yGV4774). * indicates estradiol addition. **C**. Comparison of percentage of cells with separated spindle poles with and without induction of Pch2 in *dmc1Δ rad51Δ* (yGV4774). For induction of Pch2, Estradiol was added 7 hours post-induction of meiosis (as indicated by *). **D** and **E.** Schematics explaining the model of the dual role Pch2 activity plays in both activation and silencing the G2/prophase checkpoint, in relation to ongoing chromosome synapsis. SC indicates synaptonemal complex formation between synapsing pairs of homologous chromosomes.

Overall, we here provide a framework that conceptualizes checkpoint signaling in meiotic G2/prophase and the homeostatic roles of the Pch2-Hop1 module therein (Figure 4D and E). We initially show that Pch2 is important to set a dynamic relationship between chromosomal and non-chromosomal Hop1, and Zip1-dependent synaptonemal complex assembly onto Hop1-containing meiotic chromosomes. Based on these observations, we demonstrate that the Pch2-Hop1 module is crucial for checkpoint responses to both synapsis defects and recombination failure. We posit that signaling triggered by these cellular defects share a similar signaling logic: a reliance on the presence/conversion of Hop1 into a state (*i.e.* U-Hop1) that is competent to be rapidly incorporated into chromosome-based checkpoint signaling. The function of Pch2 in the meiotic G2/prophase checkpoint can be viewed as biochemically analogous to the role of Pch2/TRIP13 in fueling mitotic spindle checkpoint function: by constantly generating the open topological state of a HORMA domain (*i.e.* Hop1 or Mad2, respectively), the AAA+ ATPase activity functions as a key catalytic engine that provides a key substrate (*i.e.* U-Hop1 or O-Mad2, respectively) available to downstream signaling machineries that generate the biochemical entities that enforce cell cycle arrest (*i.e.* chromosome-based, Mec1/Tel-activated RHMc or kinetochore-based MCC, respectively).

Paradoxically, Pch2 activity can also lead to checkpoint silencing, which again is in line with the analogy between meiotic checkpoint and mitotic spindle checkpoint signaling. We posit that both checkpoint activation and inactivation activities of Pch2 derive from the same biochemical reaction: ATP hydrolysis-driven conversion of C-Hop1 into U-Hop1. However, depending on the subcellular localization and context of the C-Hop1 client, this catalysis can lead to distinctly different outcomes: checkpoint complex disassembly and checkpoint silencing is promoted when the C-Hop1 that is converted is incorporated into an active signaling complex (RHMc in the case of the meiotic G2/prophase checkpoint). Synaptonemal complex polymerization, which drives chromosomal association of Pch2 provides a switch in ‘client’ preference during meiotic G2/prophase (Figure 4E). In the analogy with SAC regulation and MCC disassembly, SC formation can conceptually be equated with chromosome bi-orientation during mitosis; the cytological state when MCC disassembly needs to be promoted [9, 16, 52]. We note that Pch2 is likely not the only factor that contributes to meiotic checkpoint silencing (*e.g. pch2* cells eventually exit meiotic G2/prophase; *e.g.* Figure 2B and Supplementary Figure 3A), arguing for additional pathways that promote checkpoint inactivation [69], similar to what has been described for SAC/MCC inactivation [12, 70].

Despite the analogous biochemical basis of regulation between these distinct cell cycle checkpoints, we also note a fascinating distinction: Meiotic HORMA domain-containing proteins (*i.e.* Hop1 and homologs) are unique among HORMA proteins in that they encode a HORMA-interaction motif (*i.e.* closure motif, CM) in their COOH-terminal extension [10, 11, 22]. In absence of active conversion, intramolecular closed HORMA domain-CM interactions (that are predicted to generate ‘inactivated’ C-Hop1 proteins; intra-C-Hop1) seem inevitable. This implies that once Pch2 recruitment to chromosomes upon SC formation is complete, Pch2-dependent release of Hop1 from chromosomes should rapidly lead to a large pool of inactive, non-chromosomal intra-C-Hop1. We speculate that this characteristic might instate ‘switch-like’ behavior in checkpoint function downstream of SC-dependent recruitment of Pch2.

Revealing the biochemical basis of Pch2-recruitment to chromosomes, and understanding whether additional regulation drives the switch of Pch2 from ‘monomeric’ C-Hop1 to chromosome-associated C-Hop1 are questions that warrant investigation. Finally, the fact that two checkpoint cascades that respond to distinct chromosomal defects show analogous biochemical signaling logic raises fascinating questions regarding the evolutionary history and origin of these essential signaling pathways [71].

## Acknowledgements

We thank the Vader and Bird (Max Planck Institute of Molecular Physiology, Dortmund, Germany) laboratories for ideas and discussions. We thank Divya Singh (Max Planck Institute of Molecular Physiology, Dortmund, Germany) for helpful discussions on the experimental design. We thank Andrea Musacchio (Max Planck Institute of Molecular Physiology, Dortmund, Germany) for ongoing support throughout this project. We thank Adèle Marston (Welcome Trust Centre for Cell Biology, Edinburgh, UK), John Weir (Friedrich Miescher Laboratory, Tübingen, Germany), Andrea Musacchio, Divya Singh, Vaishnavi Nivsarkar, Richard Cardoso da Silva and Arnaud Rondelet (all Max Planck Institute of Molecular Physiology, Dortmund, Germany) for comments on the manuscript. We acknowledge Amy MacQueen (Wesleyan University, Middletown, CT, USA) for sharing reagents. Work in the Vader laboratory was financially supported by the European Research Council (ERC Starting Grant URDNA, agreement nr. [638197], to G.V.) and the Max Planck Society.

## Author Contribution

V.B.R. conceptualized the original idea and model of this study. V.B.R. and G.V. conceived and designed experiments. V.B.R performed all experiments and analysis. G.V. supervised the study. and G.V. wrote the manuscript.

## Declaration of Interests

The authors declare no competing interests.

## STAR Methods

### Yeast strains and constructs

Yeast strains used in this study are of SK1 background and their genotypes are listed in the supplementary data. For estradiol-dependent induction of *PCH2Δ*, a *pGAL1* promoter fusion with *PCH2Δ* was made as follows: a construct containing *pGAL1* along with a 3xFLAG epitope-6Gly-linker flanked with BglII and AscI (*BglII-pGAL-3xFLAG-6xGLY-AscI*; pGV867) restriction sites were custom synthesized into pUC57 plasmid by Genewiz Inc. The construct was recloned in a pFA6a-based precursor plasmid carrying *HIS3MX* (*pFA6a-His3MX6-PGAL1-3XFLAG-6GLY*; pGV876) Standard PCR-based one step promoter replacement strategy was employed using this construct as the template and primers 5’-TCA TAA AAA TAT TCT GAT CTC AAA CTG AAG ACA TAA AAT AAG GAT GAA TTC GAG CTC GTT TAA AC-3’ (GV2510) and 5’-AAC CCT CAG AGA TGA TCC TCG CAC TTG TAG GTC AAC TAT GTA GCT TCC ACC CCC GCC TCC-3’ (GV2511) [72].

### Growth conditions for synchronous meiosis

Yeast strains were patched onto YP-Glycerol plates and transferred to YP-Dextrose plates (containing 4% glucose). Cells were grown to saturation in liquid YPD culture at room temperature followed by inoculation in pre-sporulation media (BYTA; 50 mM sodium phthalate-buffered, 1% yeast extract, 2% tryptone and 1% acetate) at a dilution of A600 0.3. After 18 hours of growth in BYTA at 30°C, cells were washed twice in water and resuspended in sporulation media (0.3% potassium acetate) at A600 1.9 to induce meiosis at 30°C. To confirm synchronous entry into meiosis FACS analysis was used for all the experiments. For the induction experiments of *PCH2Δ*,β-estradiol (final concentration 1μM) was added at indicated time points to induce expression from the *pGAL1* promoter (note that the strains used for this also express a Gal4-ER fusion, as described [64].

### Surface spreading of chromosomes and immunofluorescence

2mL of meiotic cells were collected at indicated timepoints, 1% sodium azide was added and processed together after the last time point. Cells were treated with 500μL of 200 mM Tris pH7.5, 20 mM dithiothreitol (DTT) for 2 min at room temperature followed by spheroplasting at 30 °C in 1 M sorbitol, 2% potassium acetate, 0.13µg/µL zymolyase for 20 minutes. The spheroplasts were then gently washed two times with 1mL ice-cold MES-Sorbitol solution (1 M sorbitol, 0.1M MES pH6.4, 1 mM EDTA, 0.5 mM MgCl2) and finally resuspended in 55µL of the same solution. 20µL of the resuspended spheroplasts were placed on clean glass slides (dipped in ethanol overnight and air-dried) and two volumes of fixing solution (3% paraformaldehyde, 3.4% sucrose) added to it, followed immediately by adding four volumes of 1% Lipsol. The contents were mixed by gentle rotation of the slide. After one minute, four volumes of the fixing solution were added. A glass rod was used to mechanically spread the chromosomes. The samples were dried overnight at room temperature and stored at −20 °C. For immunofluorescence, slides were treated with 0.4% Photoflo (Kodak) diluted in PBS for 3 minutes. The slides were dipped in PBS with gentle shaking for 5 minutes. Blocking of the samples was done by incubating the samples with 5% BSA in PBS for 15 minutes at room temperature. Overnight incubation with desired primary antibodies (see below) was performed in a humidified chamber at 4 °C. The slides were subjected to two washes of 10 minutes each in PBS with gentle shaking followed by incubation with fluorescent-conjugated secondary antibody for 3 hours at room temperature. The slides were again washed twice and mounted using 20µl of Vectashield mounting media containing 4’,6-Diamidine-2’-phenylindole dihydrochloride (DAPI).

### Whole cell immunofluorescence

300 µL of meiotic cells were collected at indicated timepoints, treated with 1% sodium azide and processed together after collecting the last timepoint. Cells were harvested by centrifuging at 3000 rpm for 3 minutes and fixed overnight with 3% paraformaldehyde in 1.2 M phosphate buffer. Cells were washed 3 times with phosphate buffer and once with 1.2M sodium sorbitol. Cells were spheroplasted with 1.2M sodium sorbitol containing glusulase and zymolyase for 2 hours at 30 °C with gentle rotation. Spheroplasted cells were washed once with 1.2M sodium sorbitol solution and resuspended in 30 µL of the same solution. 5 µL of cells were allowed to settle down on a poly-L-lysine treated well of a PTFE-printed 30 well glass slide for 10 minutes. Samples were treated with ice-cold methanol for 3 minutes immediately followed by a 10 seconds treatment with ice-cold acetone. Glass slides were air-dried completely and samples incubated with 4 µL of rat -Tubulin (diluted 1:100) for 90 minutes. Samples were subsequently washed thrice with PBS followed by a one-hour incubation with 4µL of FITC-labelled secondary antibody (diluted 1:200). Samples were washed 4 times with PBS. Coverslips were mounted using 1 µL Vectashield mounting media per well containing DAPI for staining DNA. Cell cycle progression was checked by counting cells showing separated spindle poles by using tubulin stained samples. A minimum of 200 cells were counted for each timepoint.

### Microscopy and cytological analysis

Images were acquired at room temperature using 100×1.42 NA PlanApo-N objective (Olympus) on a DeltaVision imaging system (GE Healthcare) equipped with an sCMOS camera (PCO Edge 5.5). Serial z-stacks of 0.2 µm thickness were obtained and deconvolved using SoftWoRx software 6.1.l. Image quantifications for Zip1, Hop1 and Gmc2 intensity on the chromosomes were done using Imaris 7.6.4 32-bit software (Bitplane). The ‘Surface’ function was applied to the raw images on DAPI channel to identify the DNA. Values for the sum total fluorescence intensity and volume were obtained. The background was calculated using the ‘Spots’ function by manually placing three ROIs in a region that lacked DNA. Average background intensity adjusted for volume was subtracted from the chromosomal intensities of different proteins to obtain a sum of total fluorescence intensity corrected for the background. Scatter plots were generated using Graphpad Prism. Statistical significance was determined by performing Mann-Whitney U-test. For representative images, maximum intensity projection images were obtained and processed using ImageJ software.

### Western blotting

Total cell extracts were prepared by harvesting 3mL of meiotic samples at indicated time points by centrifuging at 3000 rpm for 3 minutes, stored at −20 °C and processed together. Cells were treated with 5% trichloroacetic acid for 10 minutes on ice followed by a wash using ice-cold acetone. Cells were dried overnight and lysed in 200 µL Tris-DTT buffer (Tris pH 7.4, 50mM EDTA, DTT) using glass beads on a FastPrep-24 5G instrument from MP Biomedicals. 50 µl of 5X-SDS loading buffer was added before resolving the samples by SDS-PAGE followed by antibody incubation of the samples transferred onto a nitrocellulose membrane.

### Antibodies

Chromosome surface spreads were immunostained with mouseα -GFP (Sigma-Aldrich, diluted 1:50), mouse α-Gmc2 (a kind gift from Amy MacQueen Lab, Wesleyan University, Middletown, CT, USA, diluted 1:200), goat α-Zip1 (Santa Cruz Biotechnology, diluted 1:100) and rabbit -Hαop1 (made in-house, diluted 1:200).α -Hop1 production was performed at the antibody facility of the Max-Planck-Institute of Molecular Cell Biology and Genetics (Dresden, Germany) using affinity purified full length 6-His-tagged Hop1. For whole cell immunofluorescence, rat - Tubulin (Abcam, diluted 1:100) was used. DAPI was used to stain the DNA in both cases. Following antibodies with respective dilutions were used for western blot of the protein extracts: rabbit α-Hop1 (made in-house; 1:10000), mouse α-Pgk1 (Thermo Fischer, 1:5000); rabbit α-phospho-Histone-H3-Thr11 (Abcam, 1:1000), mouse α-Flag (Sigma-Aldrich,1:1000), mouse α-Cdc5 (Medimabs, 1:1000), rabbit α-GFP (made in-house, 1:5000).

### Flow cytometry

Synchronous cell cycle progression of meiotic cultures was assessed by flow cytometry as described in [73], using an Accuri™ C6 Flow Cytometer (BD Biosciences).

## Supplementary Information

Homeostatic control of meiotic G2/prophase checkpoint function by Pch2 and Hop1

Vivek B. Raina 1,2 and Gerben Vader 1,2,*

1 Department of Mechanistic Cell Biology, Max Planck Institute of Molecular Physiology, Otto-Hahn-Strasse 11, 44227 Dortmund, Germany

2 International Max Planck Research School (IMPRS) in Chemical and Molecular Biology, Max Planck Institute of Molecular Physiology, Otto-Hahn-Strasse 11, 44227, Dortmund, Germany

## Supplementary data

Yeast strains

All strains are of the SK1 background.

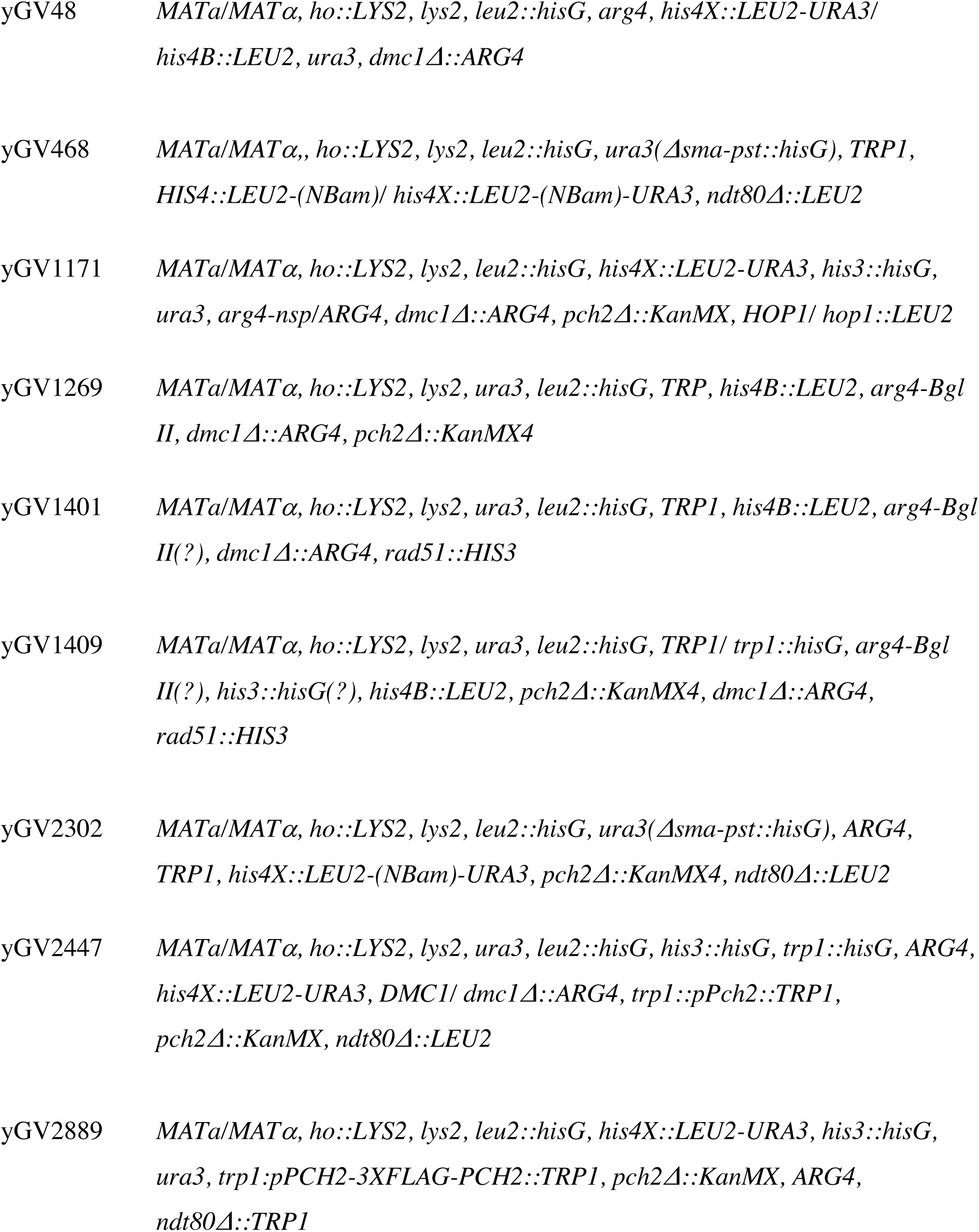
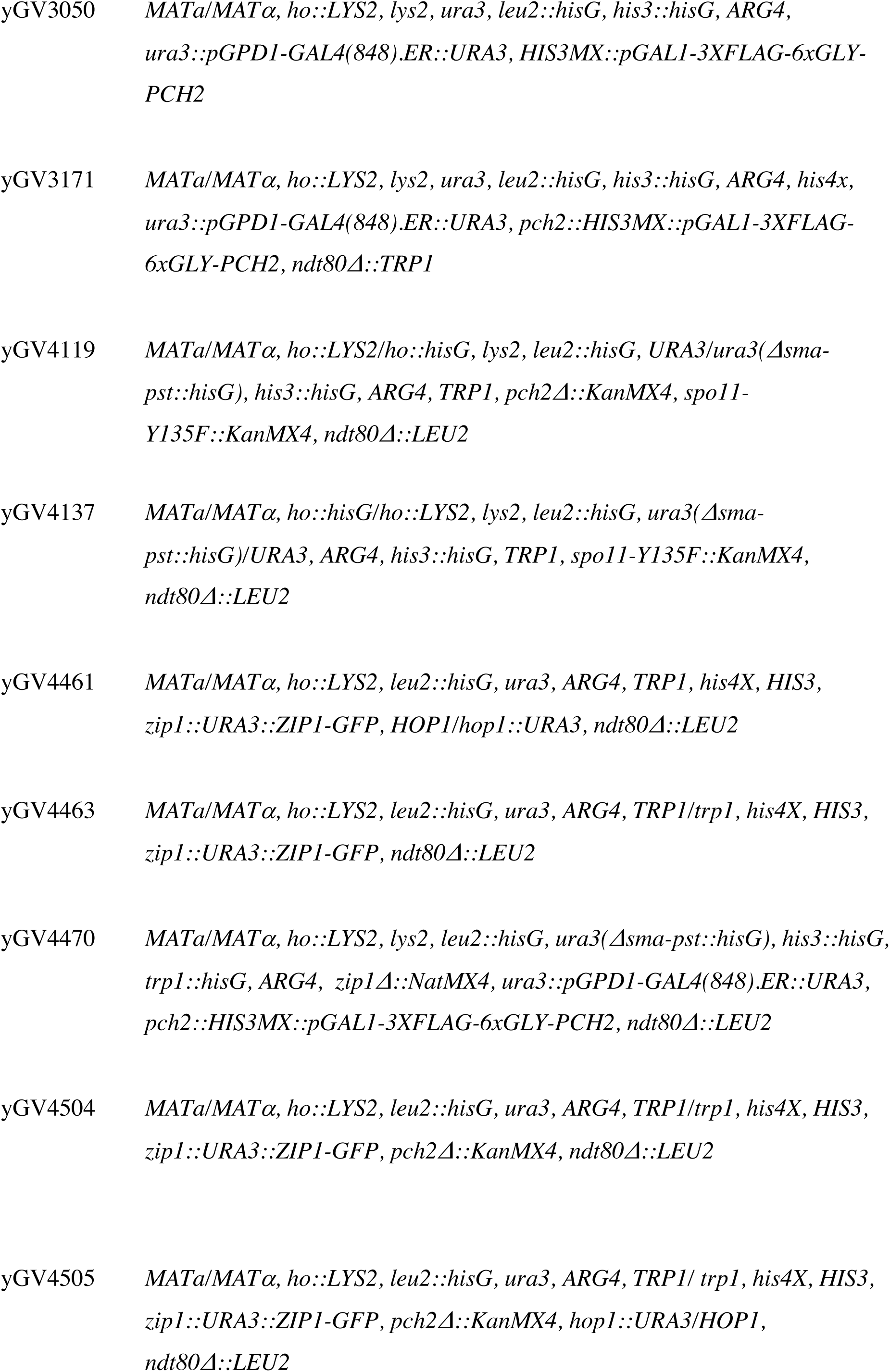
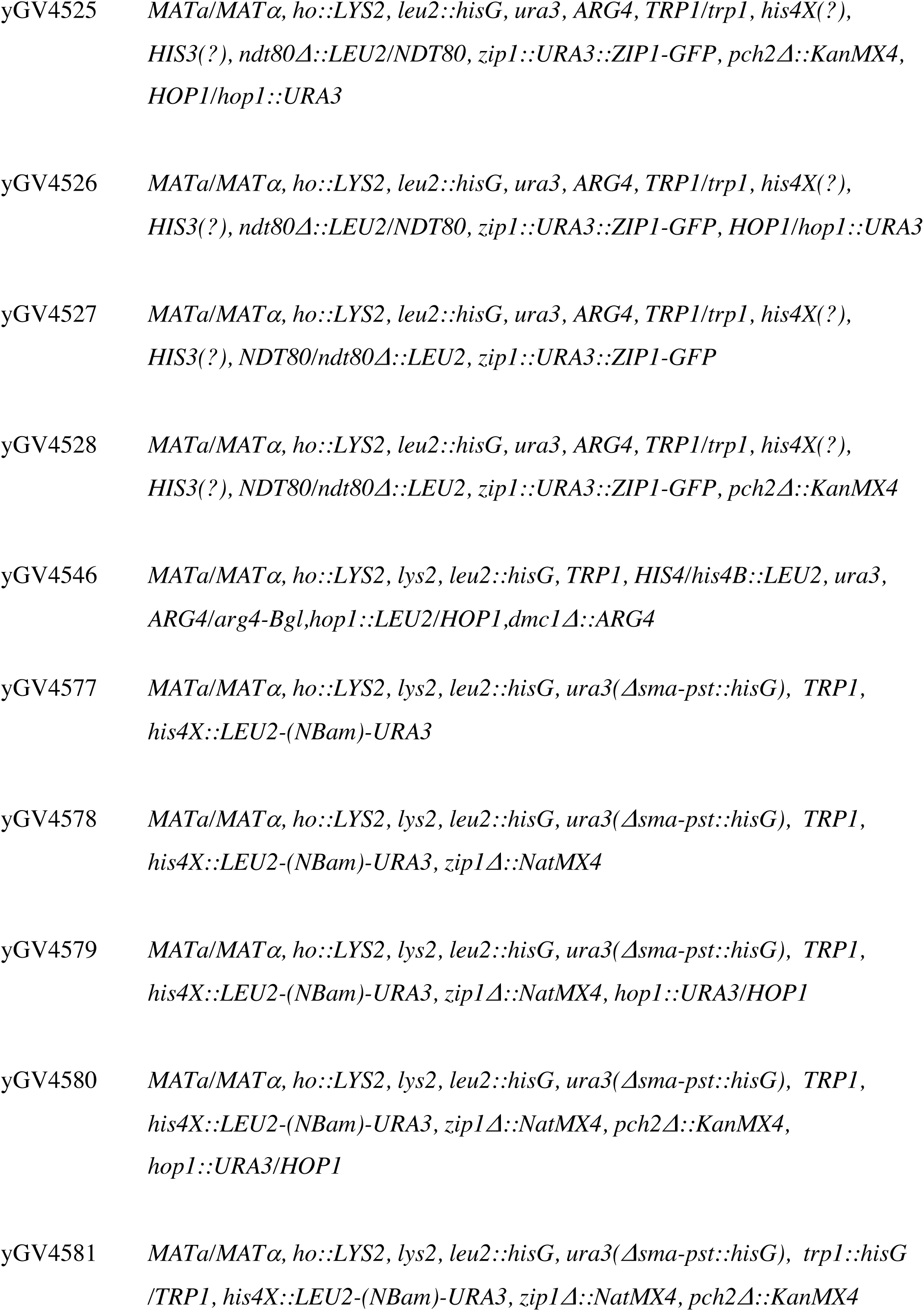
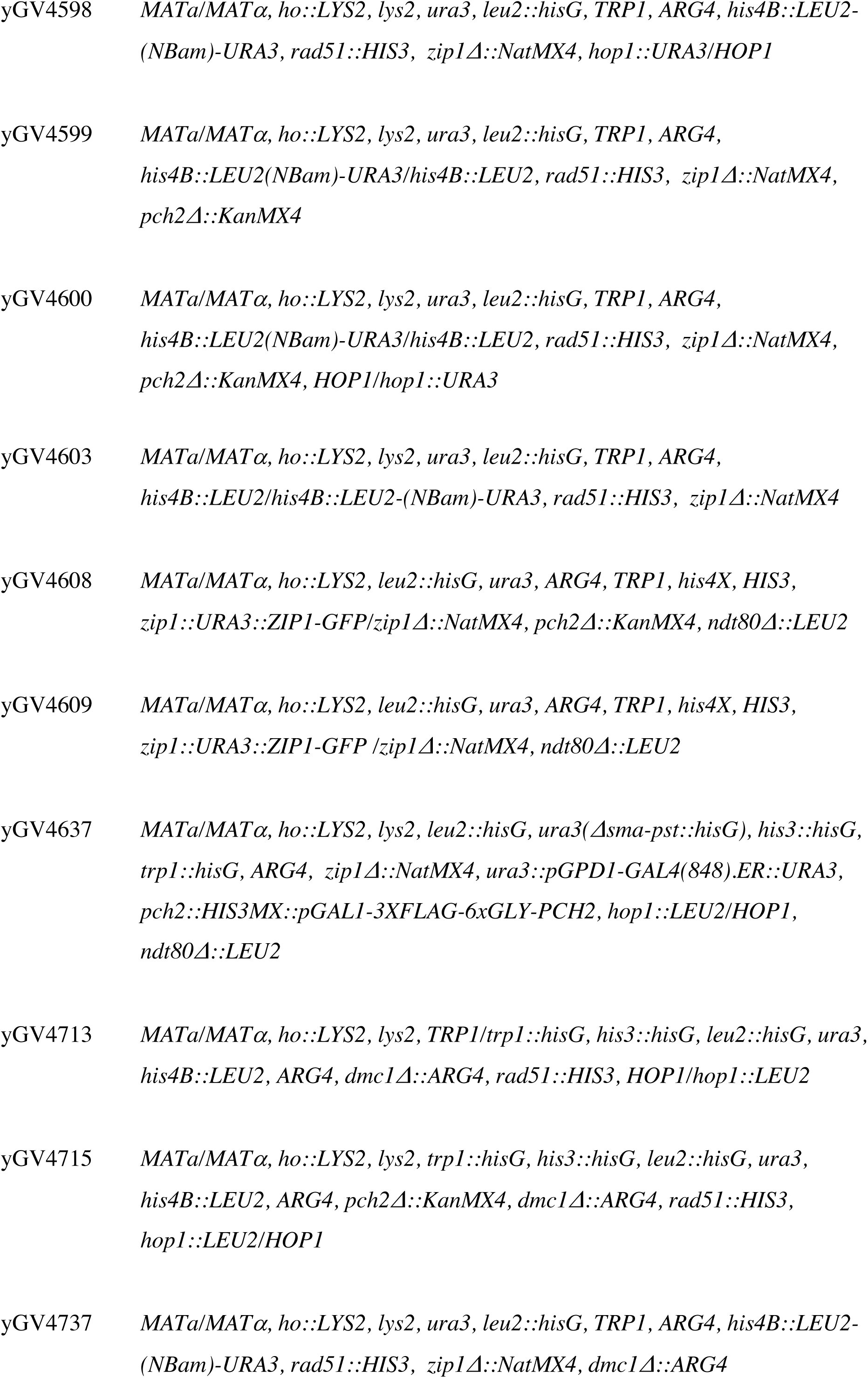
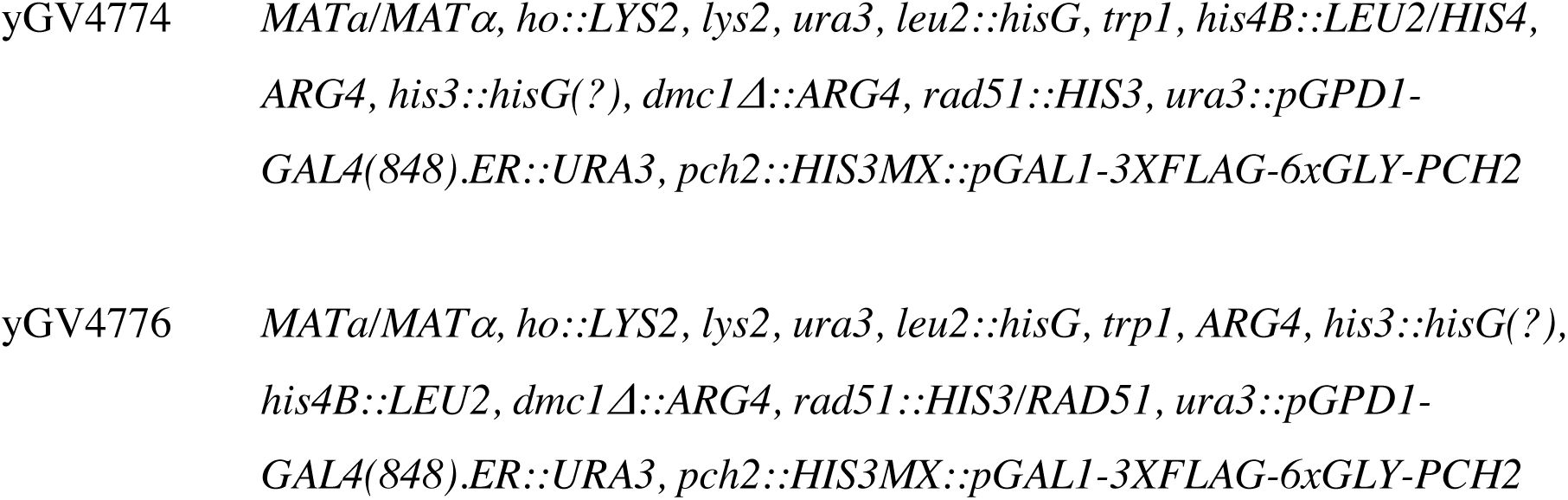

---

**Supplementary Figure 1.**
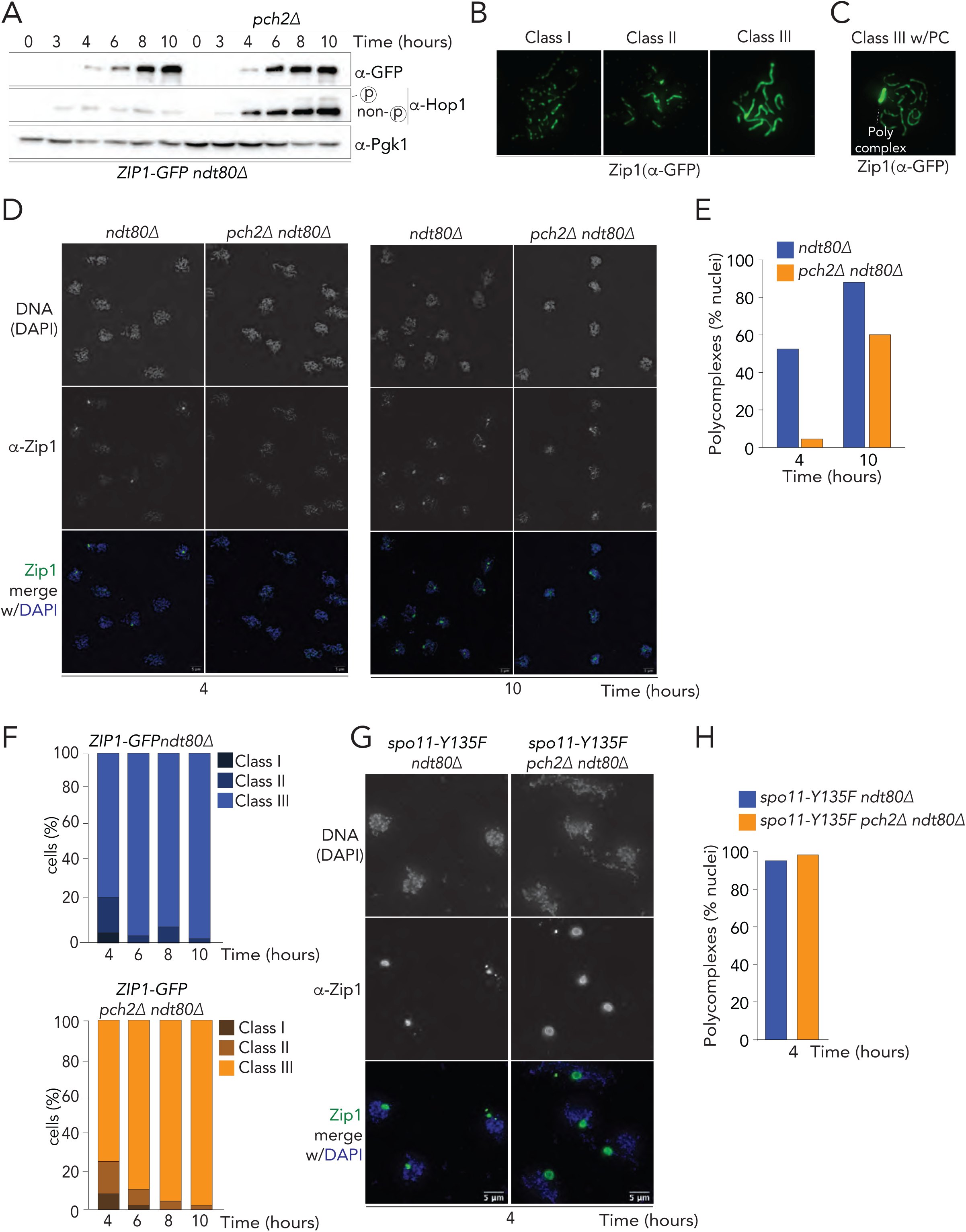
**A.** Western blot analysis of Zip1 (α-GFP) and Hop1 to compare expression levels in *ZIP1-GFP ndt80Δ* (yGV4463) and *ZIP1-GFP pch2Δ ndt80Δ* (yGV4504) at different time points. Pgk1 was used as loading control. **B.** Representative images of nuclei showing different extents of Zip1 polymerization. Class I depict foci-like appearance of Zip1, Class II depicts small stretches of Zip1 and Class III depicts extended polymerization of Zip1 (α-GFP) along chromosome lengths. Images are from *ZIP1-GFP ndt80Δ* (yGV4463) **C.** Representative image of a cell containing extended Zip1 (-GFP) polymerization (Class III) along with a PC. Images are from *ZIP1-GFP ndt80Δ* (yGV4463). PC quantifications were performed in cells exhibiting similar staining patterns. **D.** Representative images of meiotic chromosome spreads stained for Zip1 (green) using -Zip1 antibody and DNA (blue) using DAPI at indicated timepoints from *ndt80Δ* (yGV468) and *pch2Δ ndt80Δ* (yGV2302) strains. Scale bar is 5μm. **E.** Comparison of percentage of cells containing PCs at 4 and 10 hours post-induction of meiosis from the same strains as in (D). At least 100 nuclei with extended Zip1 polymerization (class III) were counted. **F.** Comparison of percentage of cells showing different extents of Zip1 polymerization divided into class I, II and III of the same strains as used in (A); *ZIP1-GFP ndt80Δ* (yGV4463) and *ZIP1-GFP pch2Δ ndt80Δ* (yGV4504) **G.** Representative images for Zip1 and DNA staining on chromosome spreads to compare effects of Pch2 on PC formation 4 hours post induction of meiosis in strains containing catalytically dead mutant of Spo11. Strains used are *spo11-Y135F ndt80Δ* (yGV4137) and *spo11-Y135F pch2Δ ndt80Δ* (yGV4119). At least 100 cells were analyzed. **H.** Comparison of percentage of cells containing PCs at 4 post induction of meiosis from the same strains as in (G). At least 100 nuclei were counted.

**Supplementary Figure 2.**
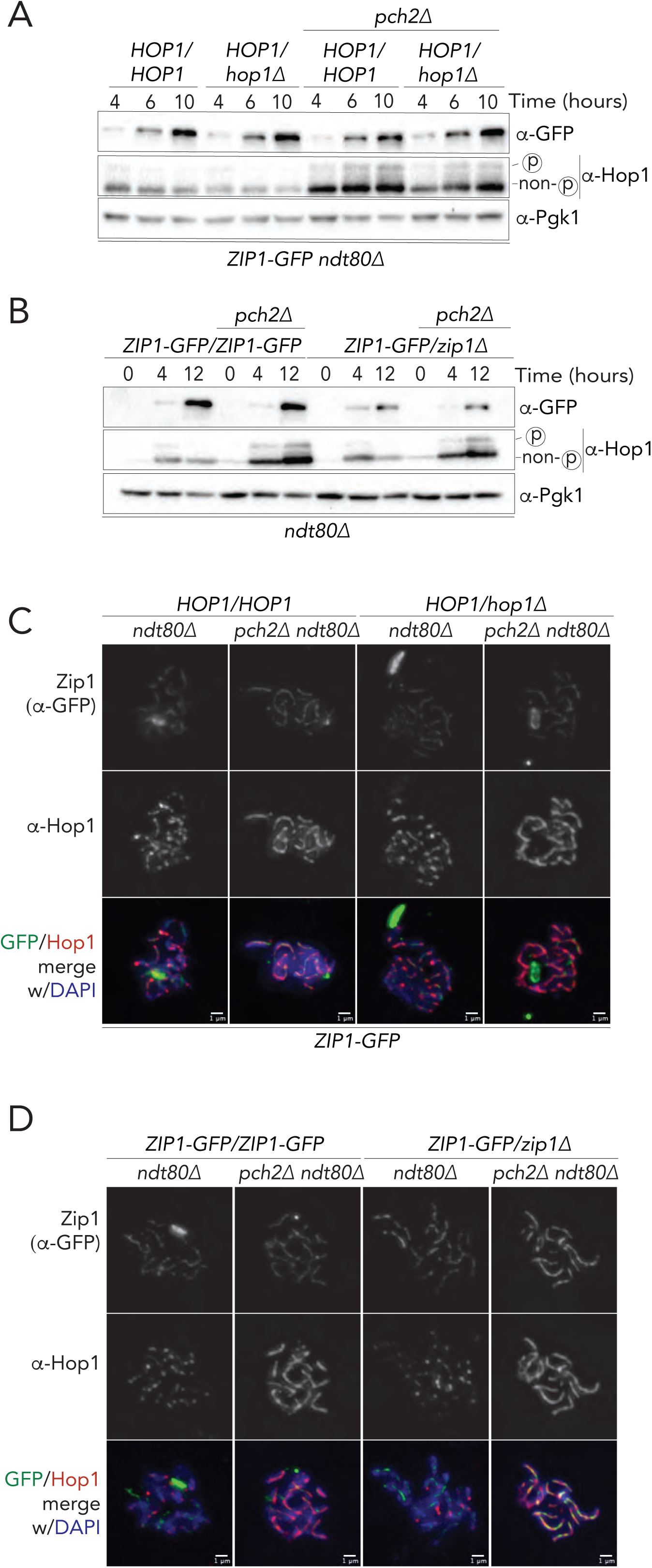
**A.** Western blot analysis to compare expression levels of Zip1 (α-GFP) and Hop1 at indicated time points from *ndt80Δ* (yGV4463), *HOP1/hop1Δ ndt80Δ* (yGV4461), *pch2Δ ndt80Δ* (yGV4504), and *HOP1/hop1 pch2Δ ndt80Δ* (yGV4505). Pgk1 was used as loading control. **B.** Expression analysis of Zip1 (α-GFP) and Hop1 at indicated time points by performing western blot of samples prepared from *ndt80Δ* (yGV4463), *pch2Δ ndt80Δ* (yGV4504), *ZIP1-GFP/zip1Δ ndt80Δ* (yGV4609) and *ZIP1-GFP/zip1Δ pch2Δ ndt80Δ* (yGV4609). Pgk1 was used as loading control. **C** and **D.** Representative immunofluorescence images showing meiotic chromosome spread of strains used in (A) and (B) respectively at 4 hours post induction of meiosis. Scale bars indicate 1µm.

**Supplementary Figure 3.**
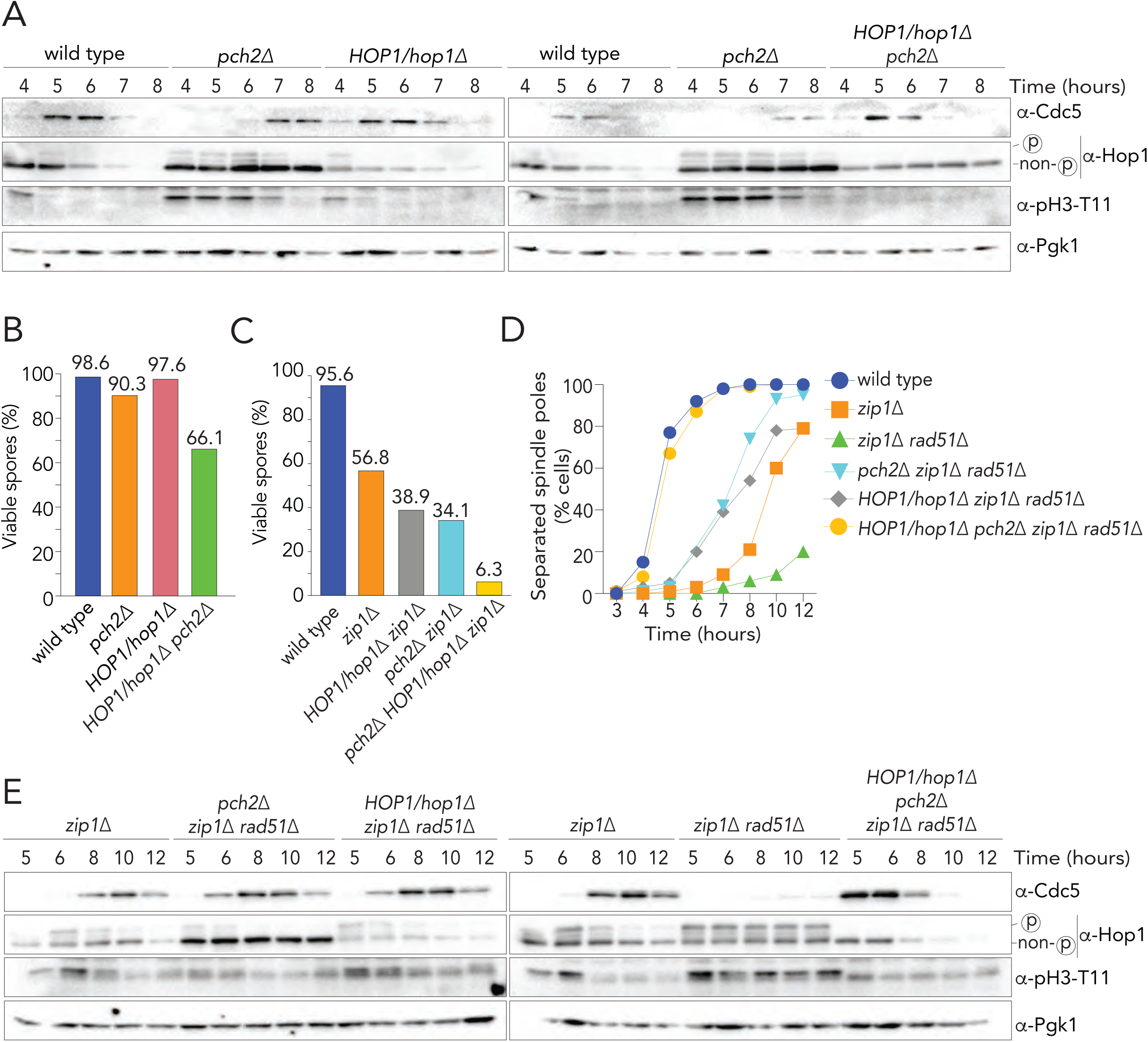
**A.**Western blot analysis to compare expression levels of Zip1 (α-GFP) and Hop1 at indicated time points from *ndt80Δ* (yGV4463), *HOP1/hop1Δ ndt80Δ* (yGV4461), *pch2Δ ndt80Δ* (yGV4504), and *HOP1/hop1 pch2Δ ndt80Δ* (yGV4505). Pgk1 was used as loading control. **B.** Expression analysis of Zip1 (-GFP) and Hop1 at indicated time points by performing western blot of samples prepared from *ndt80Δ* (yGV4463), *pch2Δ ndt80Δ* (yGV4504), *ZIP1-GFP/zip1Δ ndt80Δ* (yGV4609) and *ZIP1-GFP/zip1Δ pch2Δ ndt80Δ* (yGV4609). Pgk1 was used as loading control. **C** and **D.** Representative immunofluorescence images showing meiotic chromosome spread of strains used in (A) and (B) respectively at 4 hours post induction of meiosis. Scale bars indicate 1µm.

**Supplementary Figure 4.**
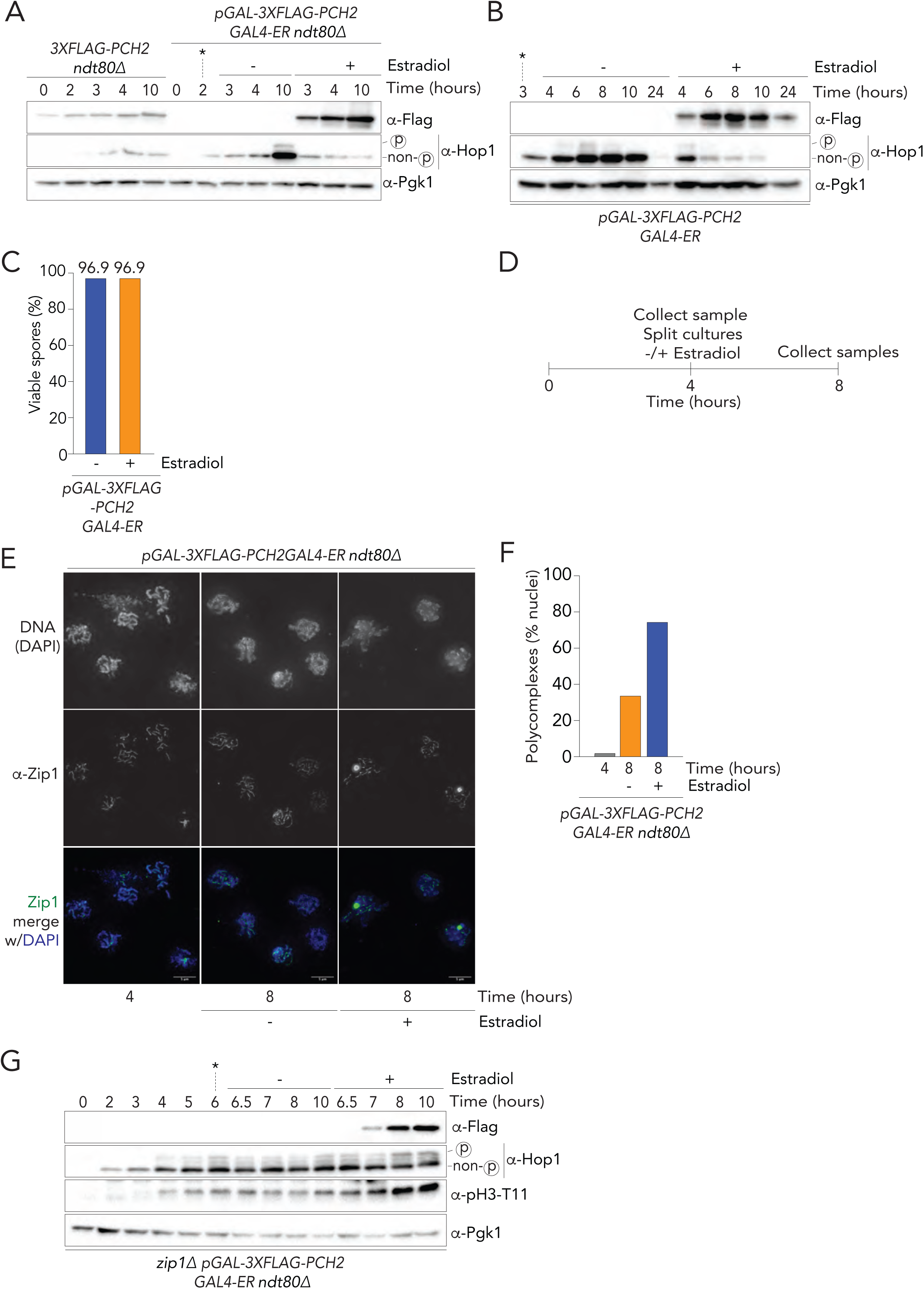
**A.** Comparison of expression levels of Pch2 (α-Flag) and Hop1 with and without induction of Pch2 in *pGAL1-3XFLAG-PCH2 ndt80Δ* (yGV3171) and *3XFLAG-PCH2 ndt80Δ* (yGV2889) strains. Pgk1 was used as loading control. Estradiol was added to induce expression of Flag-Pch2 at 2 hours post induction of meiosis (indicated by *). **B.** Expression of Flag-Pch2 was induced by estradiol addition at 3 hours post induction of meiosis (as indicated by *), and expression levels of Flag-Pch2 and Hop1 compared with and without induction at indicated time points. Pgk1 was used as loading control. Strain used is *pGAL1-3XFLAG-PCH2 (*yGV3050). **C.** Comparison of spore viability in *pGAL1-3XFLAG-PCH2 (*yGV3050) with and without induction of Pch2. Spore viability was performed from the same time course experiment as in B. See **Supplementary Table 1** for spore viability data. **D.** Schematic showing time point of addition of estradiol and time point of sample collection for the experiment shown in (E) and (F). **E.** Representative images with Zip1 (green) and DNA (blue) of meiotic chromosome spreads from *pGAL1-3XFLAG-PCH2 ndt80Δ* (yGV3171) with and without induction of 3xFlag-Pch2. **F.** Quantification of percentage of nuclei showing PCs at indicated timepoints comparing the differences with and without induction of Flag-Pch2 in the strain shown in (E). Scale bar is 5μm. A minimum of 100 cells with class III Zip1 staining were counted. **G.** Western blot analysis of Hop1 and pH3-T11 in *zip1 pGAL1-3XFLAG-PCH2 ndt80Δ* (yGV4470). Pch2 was induced by the addition of estradiol at 6 hours post induction of meiosis (as indicted by *) and expression confirmed using α-Flag. Pgk1 was used a loading control.

**Supplementary Table 1.** Spore viability data, as described in Supplementary Figure 3B, C, and Supplementary Figure 4C.

**Supplementary Table 2.** Statistical information for Figure 1C and Figure 3E.

## Notes

### Competing Interest Statement

The authors have declared no competing interest.

